# Purifying selection and phylogenetic discord among microneme proteins in *Toxoplasma gondii*

**DOI:** 10.64898/2026.03.28.714955

**Authors:** Meir Zhang, Justen Whittall, Pascale S. Guiton

**Author notes:** Author for Correspondence: JW =.

## Abstract

In *Toxoplasma gondii*, microneme proteins (MICs) are secreted components of the apical complex that play central roles in motility, host cell attachment, and invasion. Because proteins at the host–parasite interface are often predicted to evolve rapidly, MICs have been suggested as candidates for adaptive diversification. We tested this expectation using comparative analyses of three relatively understudied microneme proteins, MIC13, MIC12, and MIC16. Coding sequences were assembled from GenBank and ToxoDB, aligned by translation, and analyzed using maximum-likelihood phylogenetics, codon-based tests of selection, and predicted protein structure. MIC13 was represented by 51 sequences, MIC12 by 30, and MIC16 by 34, spanning multiple *T. gondii* haplogroups and including *Hammondia hammondi* and *Neospora caninum* as outgroups. All three genes were highly conserved among *T. gondii* strains, but their phylogenetic trees were topologically incongruent, indicating that individual MICs do not recover a single shared strain history. Contrary to expectation, no positively selected codons were detected in any gene. Instead, purifying selection was detected at one site in MIC13 and 15 sites in MIC12, while no significant codon-specific selection was detected in MIC16. Several constrained MIC12 sites overlapped annotated EGF and calcium-binding EGF-like domains, consistent with structural conservation of extracellular adhesion modules. AlphaFold prediction of MIC13 supported two sialic acid-binding micronemal adhesive repeat regions, but the single constrained MIC13 site did not overlap these motifs. Together, these results indicate that MIC13, MIC12, and MIC16 are shaped more by sequence conservation and heterogeneous gene histories than by strong recurrent positive selection. These findings refine expectations for microneme evolution in *T. gondii* and highlight conserved domains that may be important for parasite invasion and future functional study.

## Introduction

Foodborne illness is an ever-present concern in the daily lives of people globally and can impose substantial economic and public health burdens. Among these, *Toxoplasma gondii* is a major etiological agent, ranking second only to non-typhoidal *Salmonella* as the culprit in foodborne disease (Hoffmann et al., 2025). *Toxopolasma gondii* is a protozoan parasite in the Apicomplexa estimated to infect approximately 25% of the global human population (Molan et al., 2019). Although many infections are asymptomatic, chronic toxoplasmosis can persist for the lifetime of the host, and there is currently no cure for chronic infection (Hajj et al., 2021).

Treating toxoplasmosis requires a thorough understanding of the parasite and how it invades host cells, but this may be complicated if proteins involved in host–parasite interactions are rapidly evolving, as has been reported for *Plasmodium* (Gunjan & Goyal, 2025). Infection with *T. gondii* occurs primarily through ingestion of oocysts containing sporozoites from environmental reservoirs, such as water and soil contaminated with infected feline feces, with felines serving as the definitive hosts in which the parasite completes its sexual cycle (Delgado et al., 2022; Hill and Dubey, 2016). A second major route of transmission is ingestion of tissue cysts containing bradyzoites in contaminated meat from chronically infected animals. Following ingestion, parasites are released into the intestinal lumen, where they breach the intestinal mucosal barrier and actively invade epithelial host cells. Within these cells, parasites differentiate into tachyzoites inside a parasitophorous vacuole, enabling intracellular replication, egress, and dissemination throughout the host to tissues including the lungs, heart, eyes, muscles, and brain. In response to host immune pressures and other environmental cues, tachyzoites can revert to bradyzoites, establishing long-term cysts in neural and muscle tissues (Blader et al., 2015).

Because host-cell attachment, adhesion, and invasion are essential steps in this life cycle, particularly during the transition from extracellular to intracellular stages, proteins mediating these processes are central to parasite success and are often targeted for therapeutic intervention. At the same time, proteins at the host–parasite interface are frequently hypothesized to experience strong evolutionary pressures arising from host immune defenses and parasite counter-adaptations, as described in genome-scale analyses of *T. gondii* (Minot et al., 2012; Lorenzi et al., 2016). Under this framework, invasion-associated proteins are predicted to exhibit signatures of positive selection; however, their essential roles in mediating adhesion and entry into host cells may also impose strong structural and functional constraints, potentially limiting their evolutionary flexibility.

### Microneme Proteins

In order for *T. gondii* to attach to and invade host cells, it uses specialized secretory organelles called micronemes, located in the apical complex of apicomplexans (Delgado et al., 2022). These organelles secrete microneme proteins, or MICs, at the apical tip, where they are associated with parasite motility, host cell adhesion, and host cell invasion (Starnes et al. 2006; Billker et al., 2009; Dubois and Soldati-Favre, 2019) and are considered key secretory pathogenesis determinants (SPDs; Lorenzi et al. 2016). Genome-wide analyses of *T. gondii* have identified some MICs as potential targets of positive selection (Bushkin et al., 2010; Clough & Frickel, 2017; Goodswen et al., 2018) similar to the rapid evolution observed in viral envelope proteins such as the HIV env or SARS-CoV-2 spike glycoprotein.

Over 20 MICs proteins have been identified (Tagoe et al., 2021). Several MICs are well characterized (MIC2, MIC4, and MIC6), while others remain top candidates for further research like MIC16 (Wang et al. 2018), MIC13 (Ye et al., 2019), and MIC12 (Döşkaya et al., 2018). In particular, recent efforts have been made to evaluate these three less well studied MICs as potential genes for vaccine development. Immunization with a recombinant plasmid containing MIC13 has been shown to prolong survival and reduce cysts in infected mice (Yuan et al., 2013). Additionally, a recent study *in silico* modeling the structural characteristics of MIC13 has forecasted potential epitopes for recognition by B- and T-cells (Mohammadi and Dalir Ghaffari, 2025). A DNA vaccine plasmid containing MIC12 was shown to increase antibody levels and T-cell activity as well as prolong survival and reduce cyst burden in mice (Xu et al., 2025). The same effects were observed with a DNA vaccine containing MIC16 (Zhu et al., 2021). Additionally, a yeast based vaccine containing *S. cerevisiae* with surface expression of MIC16 was shown to increase antibody concentrations, increase T cell responses, and prolong survival in challenged mice (Wang et al., 2018).

Although these three MIC proteins are all localized to the microneme, they have distinct evolutionary histories. They are defined based on functional properties and are not homologous (not part of the same gene family) (Sheiner, L., et al. 2010). For example, motif analysis of MIC13 predicts two (Genbank Accession XP_002365192.1, EPT29482.1, KAF4641207.1) or three (Ye et al., 2019) sialic acid-binding micronemal adhesive repeats (MAR), which suggests they complex with this monosaccharide commonly found on the surfaces of susceptible animal cells (Varki and Schauer, 2009; Ye et al., 2019). Additionally, MIC13 was shown to contribute to *T. gondii* growth under bradyzoite-inducing stress conditions (Ye et al., 2019). MIC12 and MIC16 are both confirmed transmembrane proteins cleaved by proteases while MIC13 does not have any transmembrane domains (Sheiner et al., 2010; Mohammadi and Dalir Ghaffari, 2025). MIC12 has multiple EGF-like domains, and its specialized function is to carry a trafficking signal to the micronemes where it binds aldolase, an enzyme involved in metabolizing sugars (Sheiner et al., 2010; Liu et al., 2016). In contrast, MIC16 lacks EGF-like domains but contains thrombospondin repeats (Tsp1) (Sheiner et al., 2010).

Although no experimentally determined 3D structures exist for MIC13, MIC12, nor MIC16, the 3D structure of MIC13 has been predicted with high confidence using I-TASSER from the amino acid sequence alone (UniProt H9BC62) (Hosseininejad et al., 2023). It consists of 27.39% alpha-helices, 52.43% random coils, and 20.18% extended strands. Confidence in the 3D structure was determined using the SWISS-MODEL indicating 83.54% of residues were considered “favored” (Hosseininejad et al., 2023). Neither of the other two MICs investigated herein have previously reported 3D structures, however all three are now available on AlphaFold (Jumper et al., 2021). MIC13 has an average predicted local difference distance test score (pLDDT) of 81.88 (considered “high” confidence), whereas AlphaFold 3D models for the other two proteins are much lower (MIC12 average pLDDT = 32.84 = “very low” and MIC16 average pLDDT = 58.31 = “low”).

From an evolutionary perspective, proteins that directly mediate host–pathogen interactions are often predicted to show signatures of positive selection, reflecting adaptive responses to host defenses (Cobey, 2014). Under this expectation, microneme proteins would be anticipated to exhibit elevated rates of non-synonymous substitution relative to synonymous substitutions (i.e. positive selection) (Lorenzi et al., 2016; Yoshizaki et al., 2019). Positive selection has been previously reported for *MIC16* in a study using 12 strains by Liu et al., (2016), although no statistics such as dN-dS values were reported which begs for a more rigorous and quantitative analysis and reporting. Neither of the other two MICs (MIC12 nor MIC13) have dN-dS values reported in the literature, but this approach helps identify parasite-host interactions and the particular residues involved in parasite invasion and hosts’ natural defenses (Cobey, 2014).

### Strain Diversity

*Toxoplasma gondii* exhibits remarkable genetic diversity across its strains (Sibley et al., 2009). These strains are categorized into 16 main haplogroups, falling within six clades, each having distinct epidemiology, virulence, and evolutionary histories (Galal et al., 2019; Montoya and Liesenfeld, 2004; Khan et al., 2011; Su et al., 2012). The ME49 strain of *T. gondii* is a widely used laboratory model representing the Type II lineage, which predominates in human infections. In contrast to highly virulent Type I strains such as RH, ME49 retains the ability to form tissue cysts and establish chronic infection, making it a biologically relevant model for host–parasite interactions. Additionally, ME49 serves as the reference genome for *T. gondii*, further contributing to its widespread use in experimental and comparative genomic studies (Kissinger et al., 2003; Sibley et al., 2009; Lorenzi et al., 2016; Harb et al., 2020).

Geography plays a significant role in *T. gondii* evolutionary history with famously deep evolutionary divergence between North America and South American strains (Minot et al., 2012). Genome-wide assessments of *T. gondii* evolutionary history reveals large haploblocks nested within a subset of the 14 chromosomes (chr Ia, XI, and XII per Minot et al., 2012) which have highly conserved gene content, but are often not reflective of the evolutionary history of the rest of the genome. The reason for this is unclear, but could be due to relatively rare recombination in the wild (Grigg and Sundar, 2009) coupled with strong selection for maintaining adjacent gene content in these haploblocks (Minot et al., 2012) and/or local admixture amidst more common recombination events followed by clonal expansion of selected gene combinations (Minot et al., 2012; Lorenzi et al., 2016). The frequency and implications of sexual recombination in driving the evolutionary history of *T. gondii* is unresolved (Grigg and Sundar, 2009). Regardless, the diversity of strains and their heterogeneous evolutionary histories provide a unique opportunity to assess whether individual microneme proteins reflect shared strain histories or gene-specific evolutionary trajectories.

For comparison to *T. gondii*, many studies rely on the closely related coccidian parasites that also form tissue cysts, *Hammondia hammondi* and *Neospora caninum*. They can provide reliable outgroups that are different, yet most loci are still homologously alignable at the nucleotide and amino acid levels. In full genome comparisons, these two outgroups share 6,308-7095 OrthoMCL clusters out of the ∼7000-8000 protein coding genes detected in genome sequencing, which is more than two times that shared with the more distantly related *Sarcocystis neurona* (Lorenzi et al., 2016).

There have been several previous studies of *T. gondii* strain diversity using phylogenetic analysis of these three rarely studied MICs. Ren et al. (2012) used maximum parsimony of 18 *MIC13* sequences from *T. gondii* clinical isolates, reporting low variability for this gene and inconsistency with other genetic markers like PK1 and SAG2 (Su et al., 2006). For *MIC16*, another study used Bayesian inference and maximum parsimony of 12 sequences from clinical isolates from a diversity of mammals, constructing a tree that included clusters corresponding to the three clonal lineages (Liu et al., 2016). However, no trees built on *MIC12* sequences comparing strains were found through a literature search. Although numerous *T. gondii* complete genomes exist in Genbank and ToxoDB (Kissinger et al, 2003; Gajria et al., 2008), there are no recent reports of strain-level phylogenetic analyses and molecular evolution across multiple MICs using modern phylogenetic and molecular evolutionary methods.

### Research Questions

Here we present a bioinformatics approach investigating the relationships of *T. gondii* strains using three evolutionarily distinct microneme proteins. By comparing different MIC proteins, we can begin to understand the evolutionary history of these proteins that are involved in parasite invasion to improve our understanding of *T. gondii* pathogenesis. Additionally, by identifying sites under selection and linking them to functional domains and their position in the inferred 3D structure, we can potentially narrow the possible targets of *T. gondii* virulence. This study aims to conduct parallel bioinformatics analyses of three microneme proteins, MIC13, MIC12, and MIC16, focusing on the following specific questions. First, how do microneme proteins vary among strains of *Toxoplasma gondii*? We approach this question by inferring and comparing phylogenetic relationships among as many *T. gondii* strains using each of these three MICs. Second, are microneme coding sequences highly conserved or rapidly evolving? To address this question, we will estimate the relative rates of synonymous and non-synonymous substitutions in these MICs to determine if there are regions under purifying or positive selection. Third, do amino acid sites that exhibit evidence of significant molecular evolution overlap with known functional domains within the protein’s 3D structure? This involves cross referencing known functional domains and tertiary structure with amino acid positions undergoing positive or negative selection. We predict that different microneme proteins experience different evolutionary constraints across strains, reflecting distinct functional roles during host invasion and parasite survival. Because microneme proteins directly mediate host cell invasion, they are often assumed to experience rapid evolution driven by host–parasite interactions; however, whether this expectation holds uniformly across distinct MIC proteins remains to be determined.

## Methods

### Finding the Reference Sequence

In order to find a reference sequence of *MIC13* from *T. gondii* for further analysis, a Genbank Entrez search was conducted with the search string “MIC13 AND Toxoplasma[organism].” A RefSeq sequence was available from the canonical laboratory model strain ME49, and this became our *MIC13* reference sequence Genbank accession number <u>XM_002365151.1</u>) The same process was used to select reference sequences for *MIC12* (<u>XM_002368795.2</u>) and *MIC16* (<u>XM_002368347.2</u>). We report results for *MIC13* first since it has the most samples and most reliable 3D structure, then for *MIC12* and *MIC16*.

### Finding homologous sequences using Basic Local Alignment Search Tool (BLAST)

To compile a list of homologous *MIC13* sequences to be aligned in Geneious Prime (v. 2025.0.3), we conducted a BLAST search with the reference sequence as the query searching the core nucleotide database (core_nt) and using the BLASTN algorithm to maximize the number and diversity of strains returned. Default parameters were used. BLAST hits consisting of sequences from a diversity of *T. gondii* strains and *N. caninum* as an apicomplexan outgroup (Yoshizaki et al., 2019) were downloaded as Genbank Complete sequence and uploaded to Geneious Prime. The same workflow was used to generate alignments for *MIC16*.

For *MIC12*, this workflow did not produce sufficient number of BLAST hits using either BLASTNnor discontiguous MegaBLAST algorithms. Instead we retrieved primarily hits with low query coverage (<10%) from non-*MIC12* loci. Therefore, we blasted all available *T. gondii* genomes for *MIC12* using “NCBI Datasets = Genome”. We limited our results to those genomes with a submitter tag indicating that annotations were available in the genome sequence. These modified BLAST searches for *MIC12* used the MegaBLASTalgorithm with default parameters. This modified workflow was then repeated for *MIC13* and *MIC16* to obtain even more strain sequences for comparison. Finally, we searched for all full-length CDSs listed in ToxoDB (Harb et al., 2020; Alvarez-Jarreta et al., 2024) as *MIC13* orthologs and paralogs, and those with at least 70% identity to the reference sequence were added. This process was repeated for *MIC12* and *MIC16*.

### Multiple sequence alignment

In order to compare the BLAST hits to the reference sequence, a multiple sequence alignment was created in Geneious Prime. The coding sequence (CDS) was extracted from the annotations of each sequence, and all CDSs were aligned using translation align to preserve the reading frames (inserting gaps in multiples of three nucleotides at a time). The translation alignment used all default settings including the Blosum62 cost matrix with “build tree using alignment” enabled. Sequences were removed if they had <70% identity to the reference sequence (not homologous) or if they were missing their start or stop codons.

### Phylogenetic Analysis

To infer the relationships between strains of *T. gondii*, phylogenetic trees of the *MIC13*, *MIC12*, and *MIC16* alignments were constructed using a maximum likelihood-based algorithm. These trees were created using the RAxML (v. 8.2.11) plugin implementing the GTR CAT I nucleotide model, activated rapid bootstrapping, and then searched for the best-scoring ML tree. The GTR CAT I model uses the generalized time reversible rate matrix to analyze substitutions while accounting for unequal rates of substitution (GTR), rate heterogeneity (CAT), and proportion of invariant sites (I). Bootstrapping was performed for all trees with 1000 replicates. To compare the relationships between strains across the different MIC genes, pruned trees were then created by hand in Mesquite (v. 3.81) to include only strains that overlapped in the alignments of all three MICs.

### Molecular Evolution

In order to test for selection acting on the three microneme proteins, we used the Single-Likelihood Ancestor Counting (SLAC) and the Adaptive Branch-Site Random Effects Likelihood (aBSREL) algorithms in Datamonkey (Weaver et al. 2018). All alignments had outgroup sequences and stop codons removed and were then stripped of gaps common across all remaining sequences using the masking tool within Geneious Prime and exported as FASTA files for Datamonkey analyses. The universal genetic code was used for all analyses, and a *p*-value of < 0.05 was used to determine significance for codons undergoing selection. For analysis, regions listed under Features in the Genbank flatfile were annotated on the dN-dS output graphs. Due to the abundance of features annotated in the MIC12 reference flatfile, only those that overlapped with sites under significant selection were annotated on the dN-dS graph. For MIC12, annotations labeled “PspC” likely reflect low-complexity or misannotated domains rather than true homology to bacterial pneumococcal proteins which would be highly unusual in this eukaryotic apicomplexan.

### 3D Structures

To look for a MIC13 3D structure to help interpret the molecular evolution results, we searched the NCBI Structures database with the search string “MIC13 AND Toxoplasma[organism]”, used our Genbank reference protein sequence as the query for a BLASTP search, and set Protein Data Bank (PDB) as the database. No PDB structure was available for MIC13, so to visualize the 3D structure, we searched the AlphaFold database using the translated protein sequence. The UniProt ID# of the result with the highest pLDDT score (H9BC57) was entered into the iCn3D viewer to locate the sites of interest. For comparison to the outgroup (*N. caninum*), which also had no PDB structure available, the above process was repeated to find an AlphaFold 3D structure of MIC13 from *N. caninum* (UniProt ID# F0VGP9), and the two AlphaFold models were aligned by Vector Alignment Search Tool (<u>VAST</u>; Madej et al., 2013) to confirm homology between the 3D structures of the proteins. A similar approach was used to search for 3D structures of MIC12 and MIC16, but no empirical PDB structures were available and the AlphaFold models were highly uncertain (low pLDDT scores - see previously reported values in Introduction).

## Results

### Finding the Reference Sequence

The three reference CDSs differed substantially in length, with MIC12 much longer than MIC13 and MIC16 (Table 1).

**Table 1.** Summary of *T. gondii MIC13*, *MIC12*, and *MIC16* mRNA reference sequence lengths from the ME49 strain.

| Gene | Genbank Accession Number | Total Sequence Length (bp)<sup>1</sup> | CDS Length<sup>2</sup> (bp) | Translated Protein Length (aa)<sup>3</sup> |
| --- | --- | --- | --- | --- |
| *MIC13* | XM_002365151.1 | 1655 | 1407 | 468 |
| ***MIC12*** | XM_002368795.2 | 7731 | 6549 | 2182 |
| ***MIC16*** | XM_002368347.2 | 3049 | 2007 | 668 |

<sup>1</sup>bp = base pairs
<sup>2</sup>includes STOP codon
<sup>3</sup>aa = amino acids

### BLAST and Multiple Sequence Alignment

Combining the reference sequence and BLAST hits for *MIC13*, we found 51 sequences with an average CDS length of 1403 ± 2.96 bp (Table 2). Of these sequences, 47 are from *Toxoplasma gondii* representing 30 different strains, one is from *H. hammondi*, and three are from *N. caninum*. The CDS alignment for *MIC13* is 1407 bp long including a 108 bp deletion in two of the three *N. caninum* sequences stretching from 658-765 bp in the alignment with no gaps present in any of the other sequences (Figure 1). Sequences within the ingroup (*T. gondii*) show >99% global pairwise identity to the reference sequence (Table 2), and the average global nucleotide pairwise identity for the alignment is 96.3%, indicating homology across the alignment (Figure 1).

**Figure 1.**
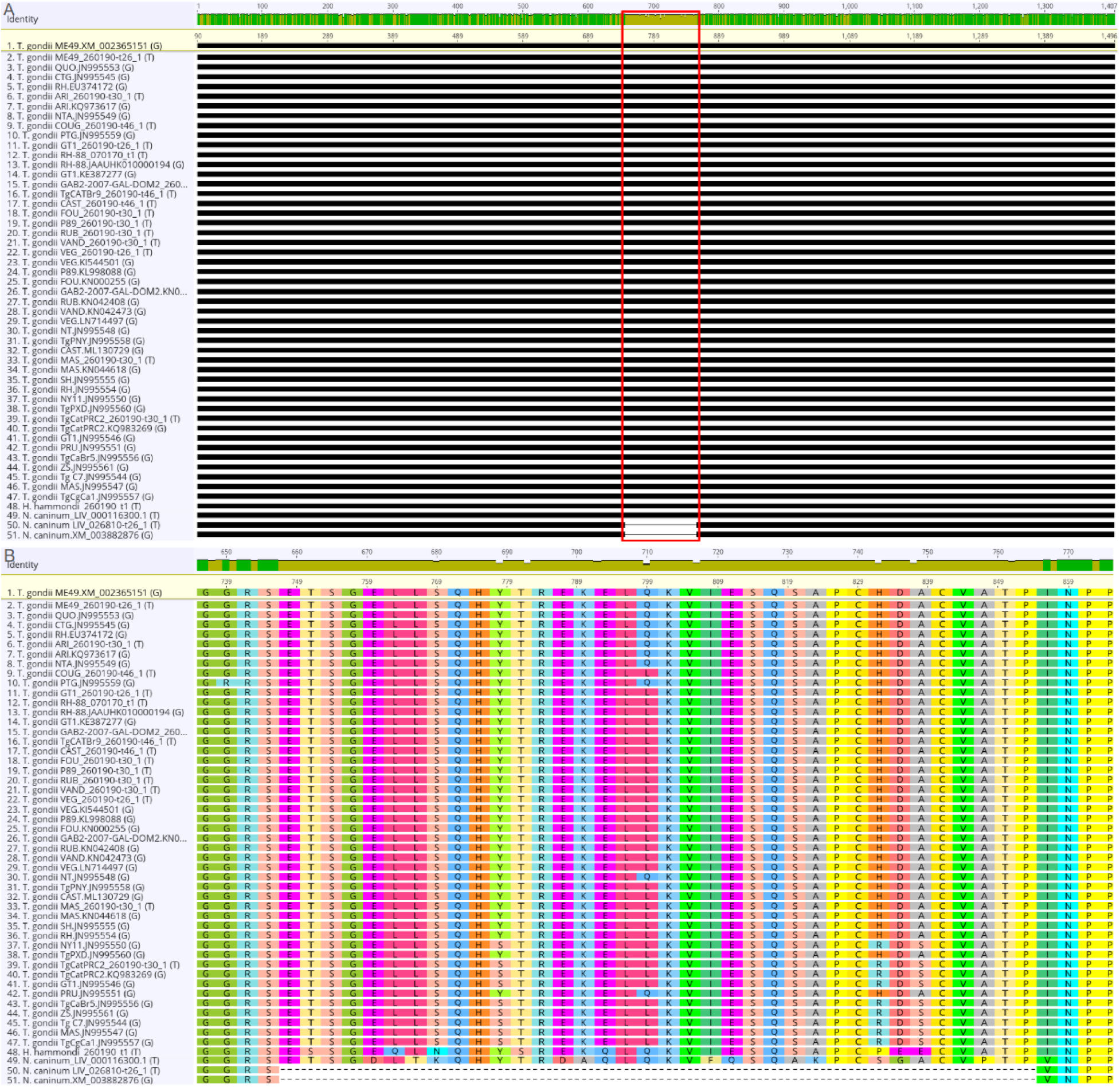
Multiple sequence alignment of *MIC13* coding sequences between *Toxoplasma gondii* strains and *Neospora caninum*. Sequences are labeled with species (and strain for *T. gondii*) followed by Genbank accession number or gene number for ToxoDB sequences. *N. caninum* and *H. hammondi* are the outgroups. This translation alignment is sorted by the number of differences to the reference sequence (*T. gondii* ME49.XM_002365151, yellow highlight). (A) shows the full CDS alignment, (B) shows the amino acids aligned from the boxed region from (A) which includes the 108 bp deletion in two of three *Neospora* sequences.

**Table 2.**
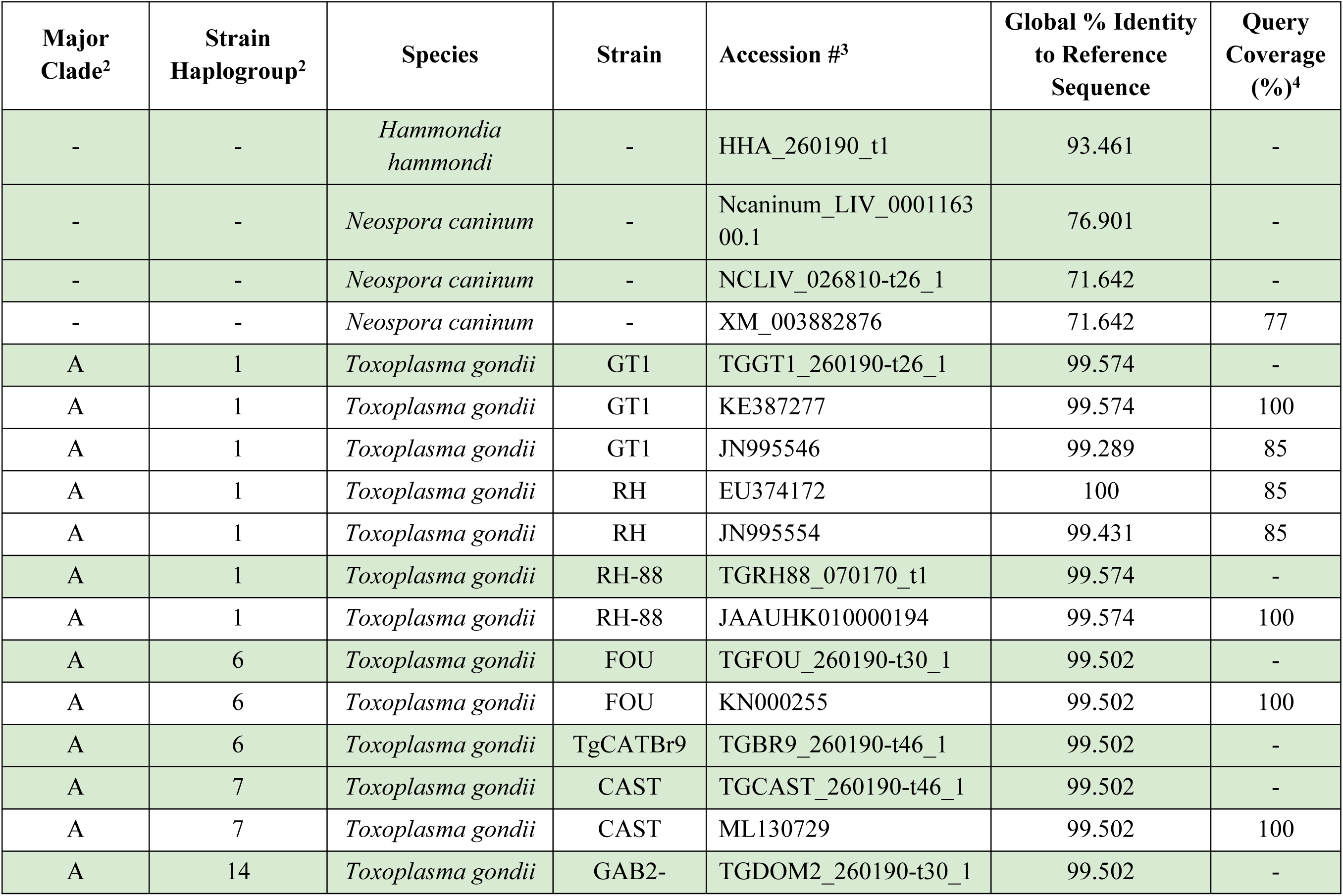
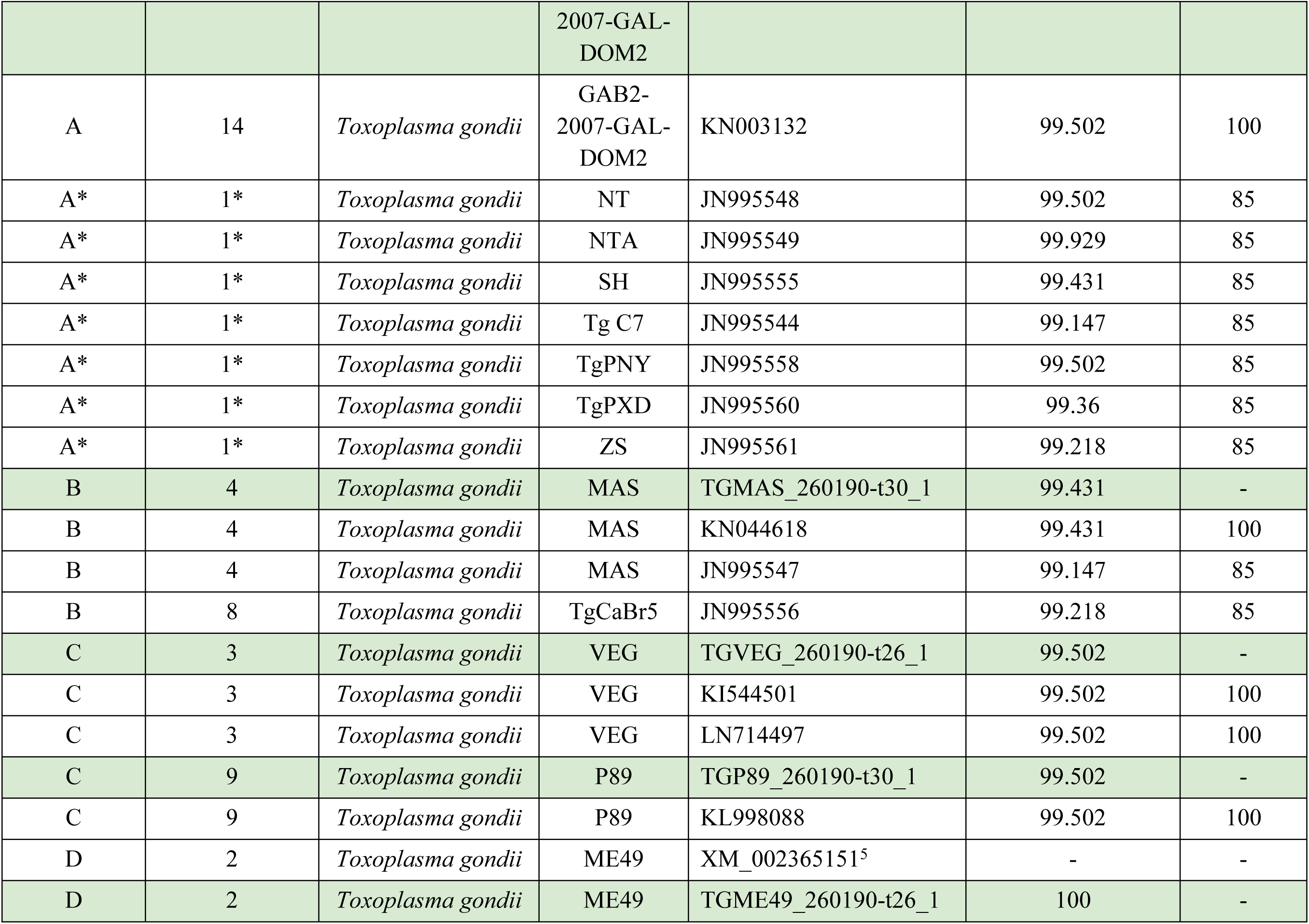
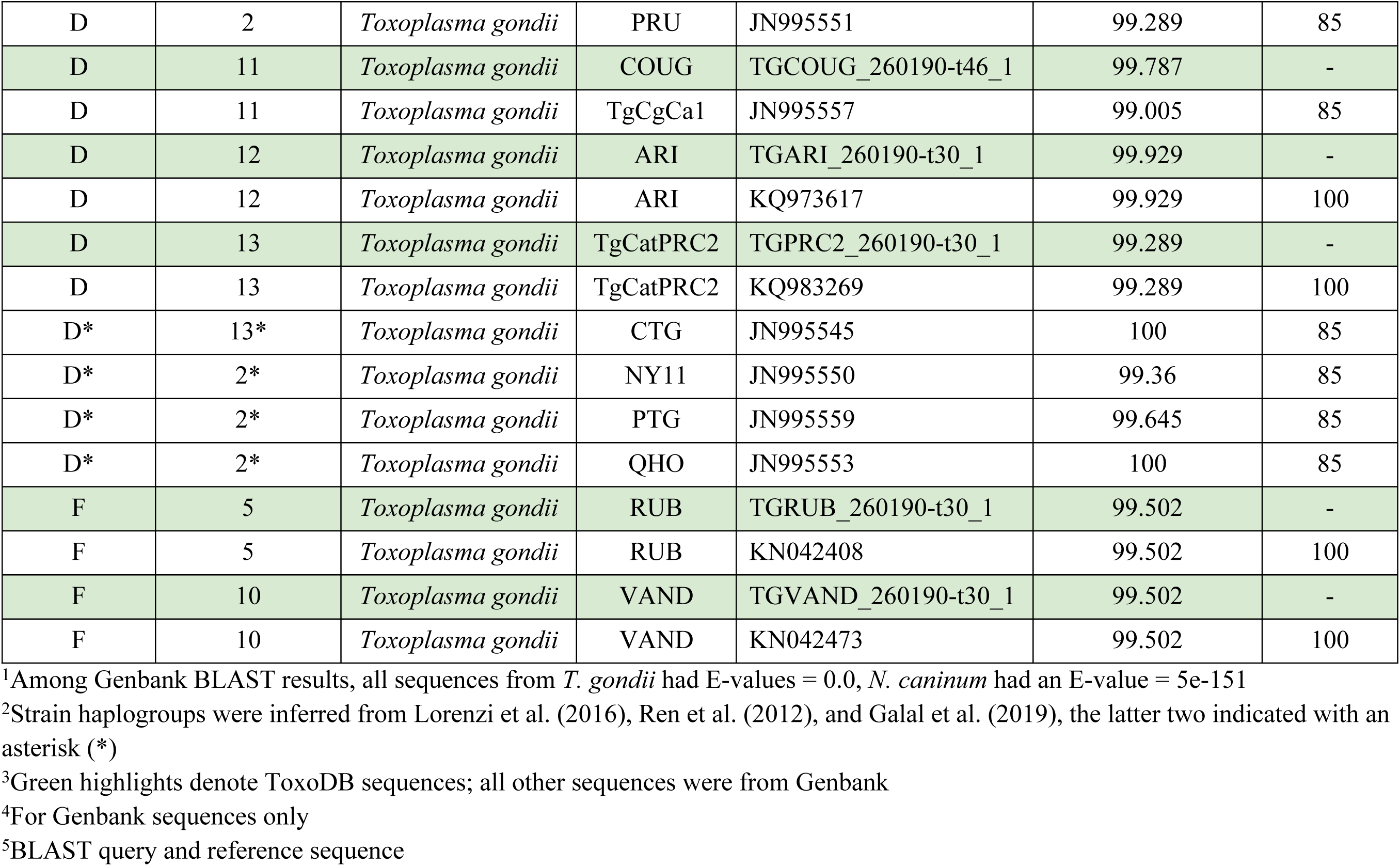
*MIC13* coding sequences from *Toxoplasma gondii* and outgroups.^1^.

A total of 30 sequences of *MIC12* were found with an average CDS length of 6369 ± 59.4 bp (Table 3). Of the sequences, 26 are from *T. gondii* representing 15 different strains, one is from *H. hammondi*, and three are from *N. caninum*. The translation alignment is 6825 bp long with an average global nucleotide pairwise identity of 93.1% (Figure 2). The *H. hammondi* sequence terminates early at alignment base pair 5406, the *T. gondii* P89 ends at base pair 5730, the *T. gondii* MAS ends at base pair 5778, and the *T. gondii* COUG ends at base pair 5883. The other *T. gondii* sequences are very similar to each other over the first 5730 base pairs, after which indels begin to separate the strains (Figure 2).

**Figure 2.**
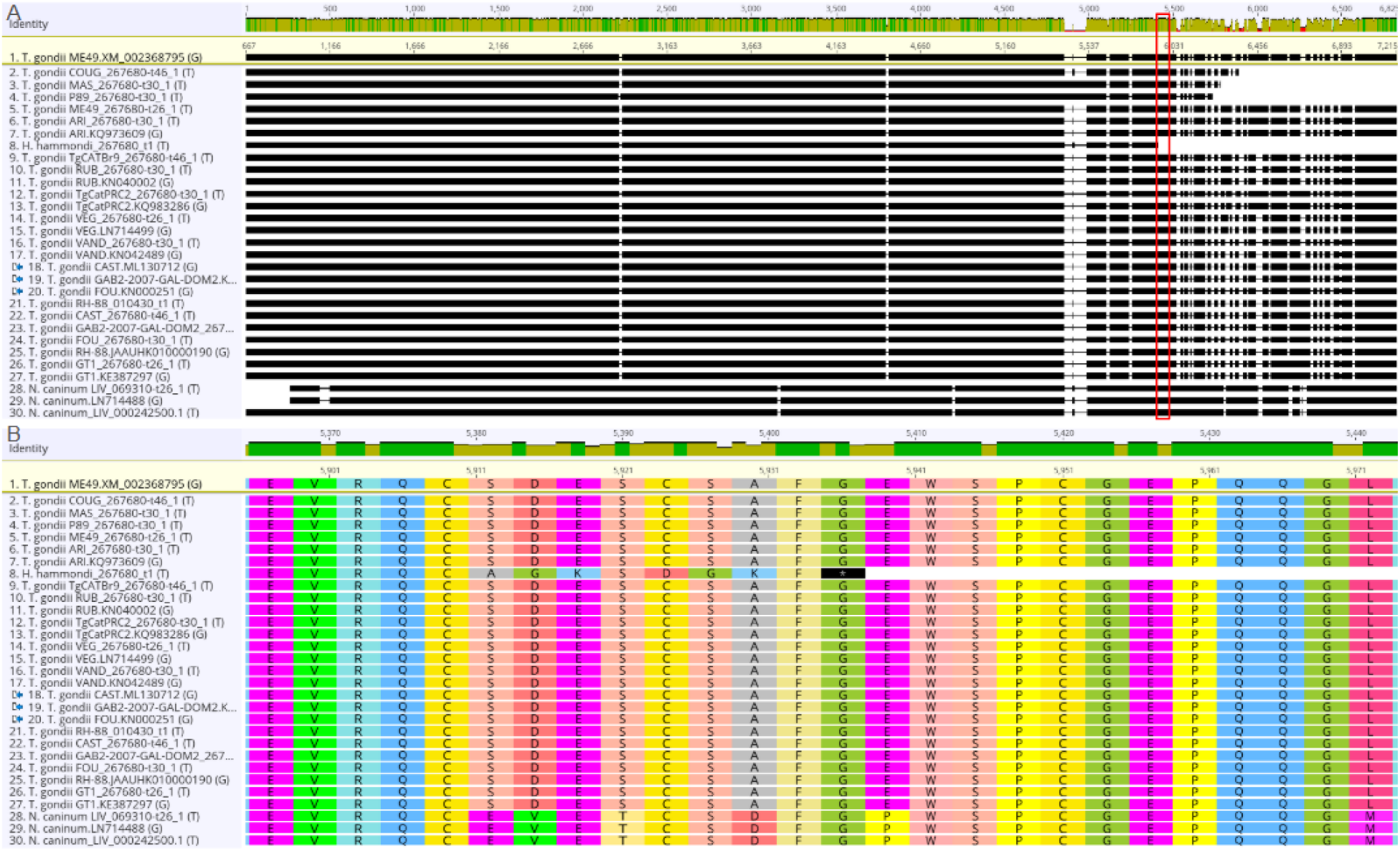
Multiple sequence alignment of *MIC12* coding sequences among *Toxoplasma gondii* strains. Sequences are labeled with species (and strain for *T. gondii*) followed by Genbank accession number or gene number for ToxoDB sequences. *N. caninum* and *H. hammondi* are the outgroups. Translation alignment is sorted by the number of differences to the reference sequence (*T. gondii* ME49.XM_002368795, yellow highlight). (A) shows full CDS alignment, (B) shows the amino acids aligned from the boxed region from (A) which includes the early stop codon in *H. hammondi.*

**Table 3.**
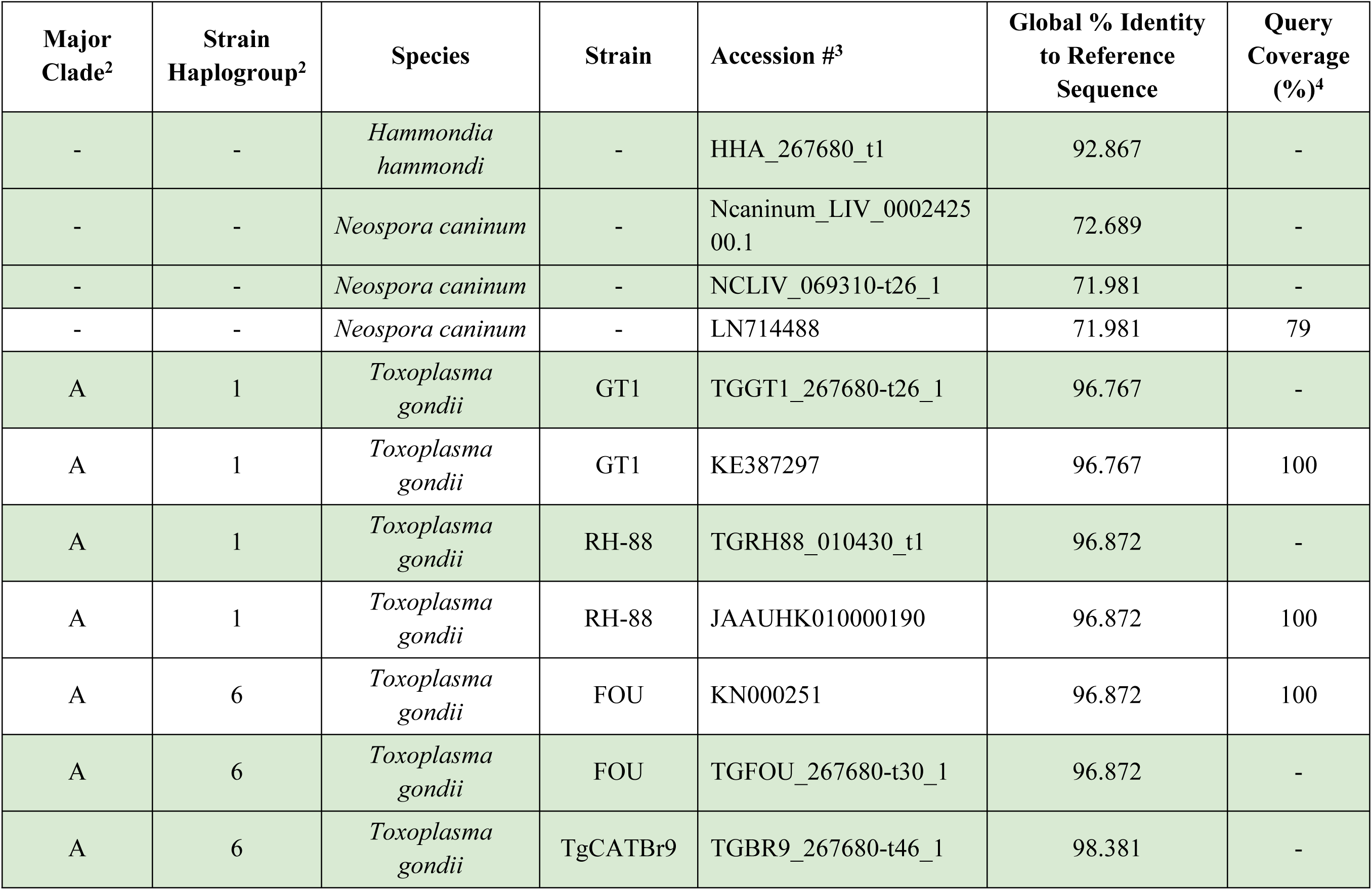
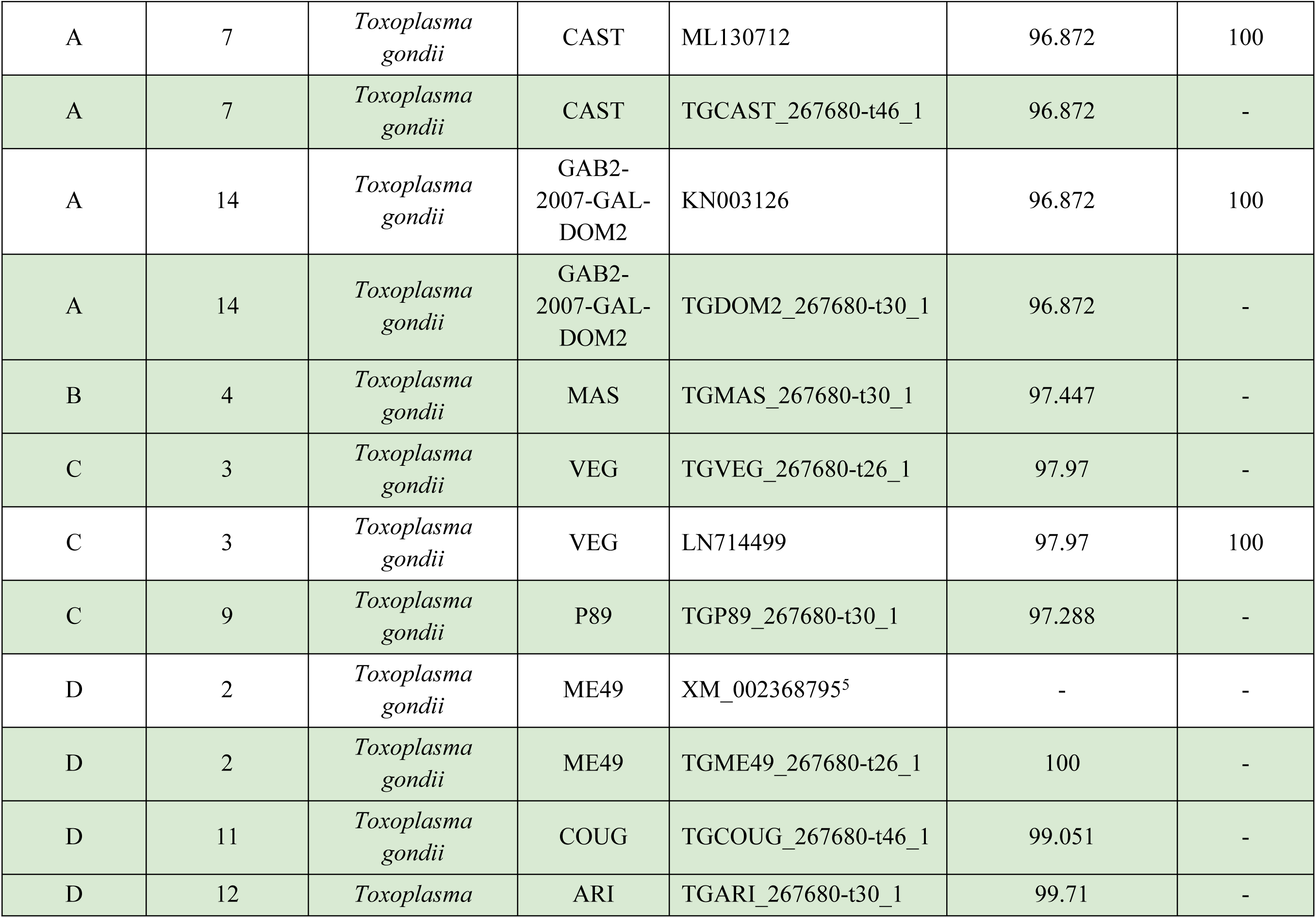
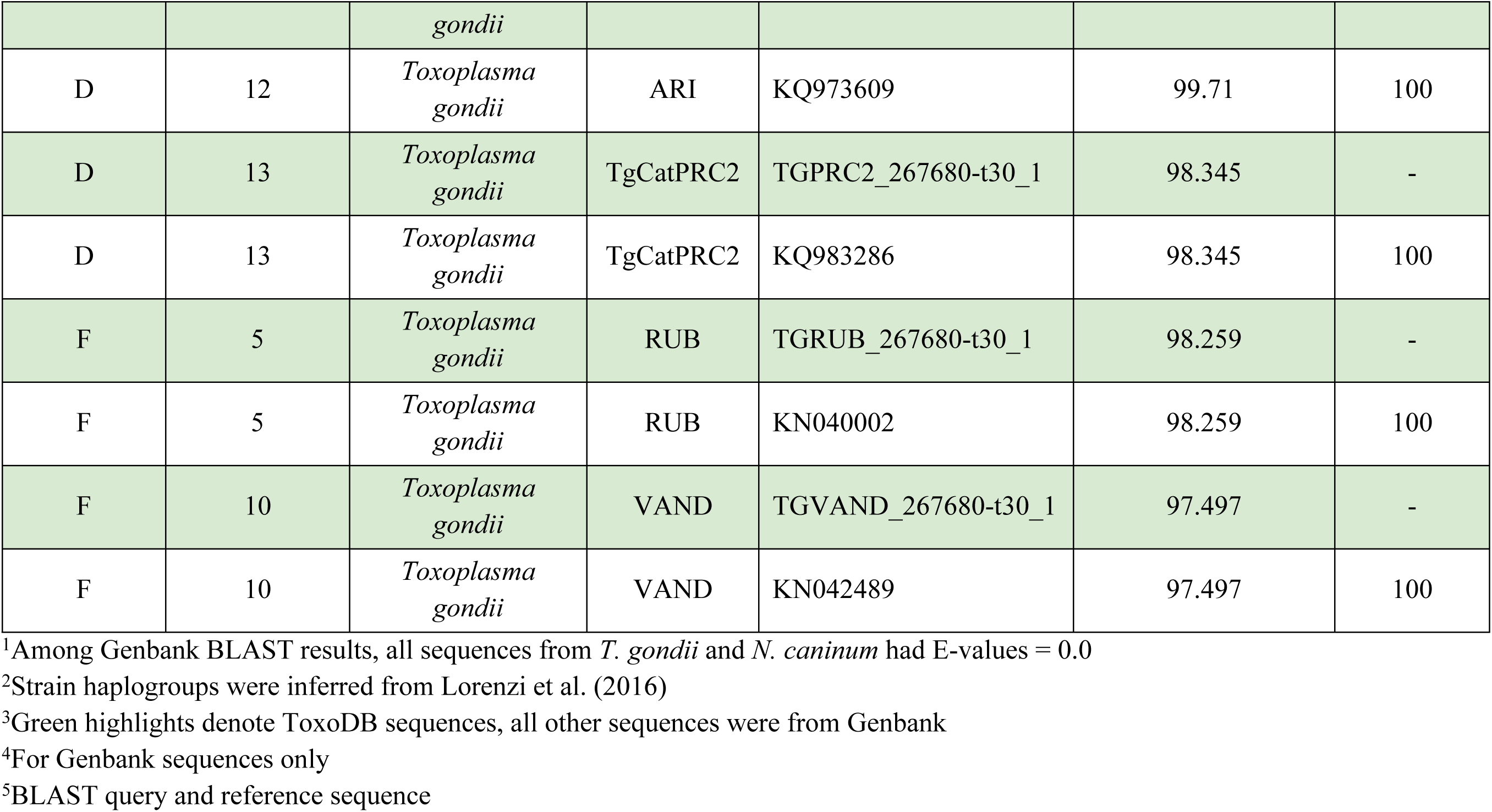
*MIC12* coding sequences from *Toxoplasma gondii* and outgroups^1^.

A total of 34 sequences of *MIC16* were found with an average CDS length of 1989 ± 9.66 bp (Table 4). Of the sequences, 31 are from *T. gondii* representing 16 different strains, one is from *H. hammondi*, and two are from *N. caninum*. The translation alignment is 2007 bp long with an average global nucleotide pairwise identity of 97.3% (Figure 3). Sequences from *T. gondii* VEG, *H. hammondi*, and *N. caninum* start later in the alignment than the other three sequences by 109 bp for *T. gondii* VEG and *H. hammondi*, 103 bp for one *N. caninum* sequence, and 286 for the other (Figure 3).

**Figure 3.**
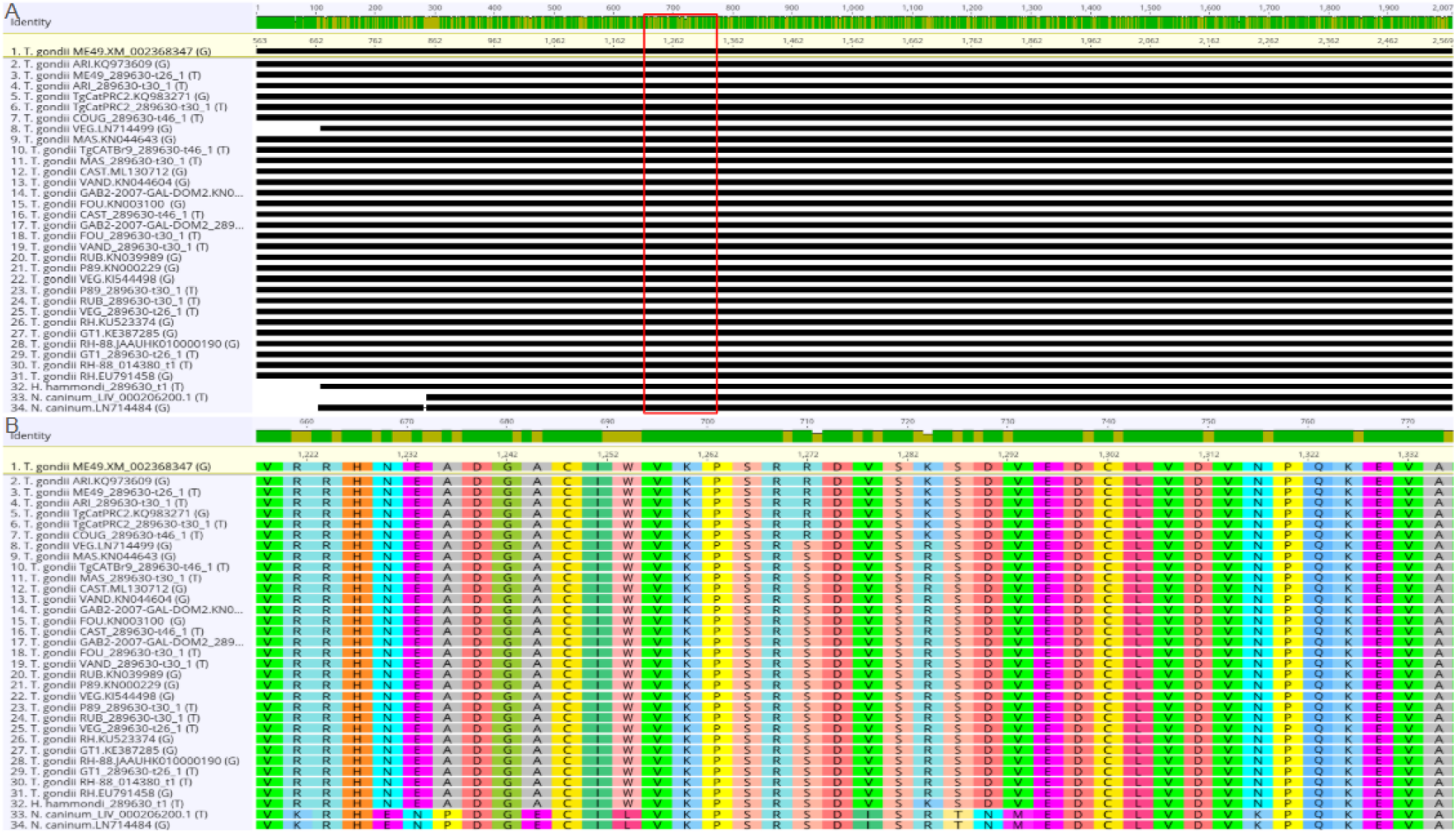
Homology of the *MIC16* coding sequences between *Toxoplasma gondii* strains and Neospora caninum. Sequences are labeled with species (and strain for *T. gondii*) followed by Genbank accession number or gene number for ToxoDB sequences. *N. caninum* and *H. hammondi* are the outgroups. Translation alignment is sorted by # differences to reference sequence (*T.gondii* ME49.XM_002368347, yellow highlight). (A) shows full CDS alignment, (B) shows translation in boxed region from (A).

**Table 4.**
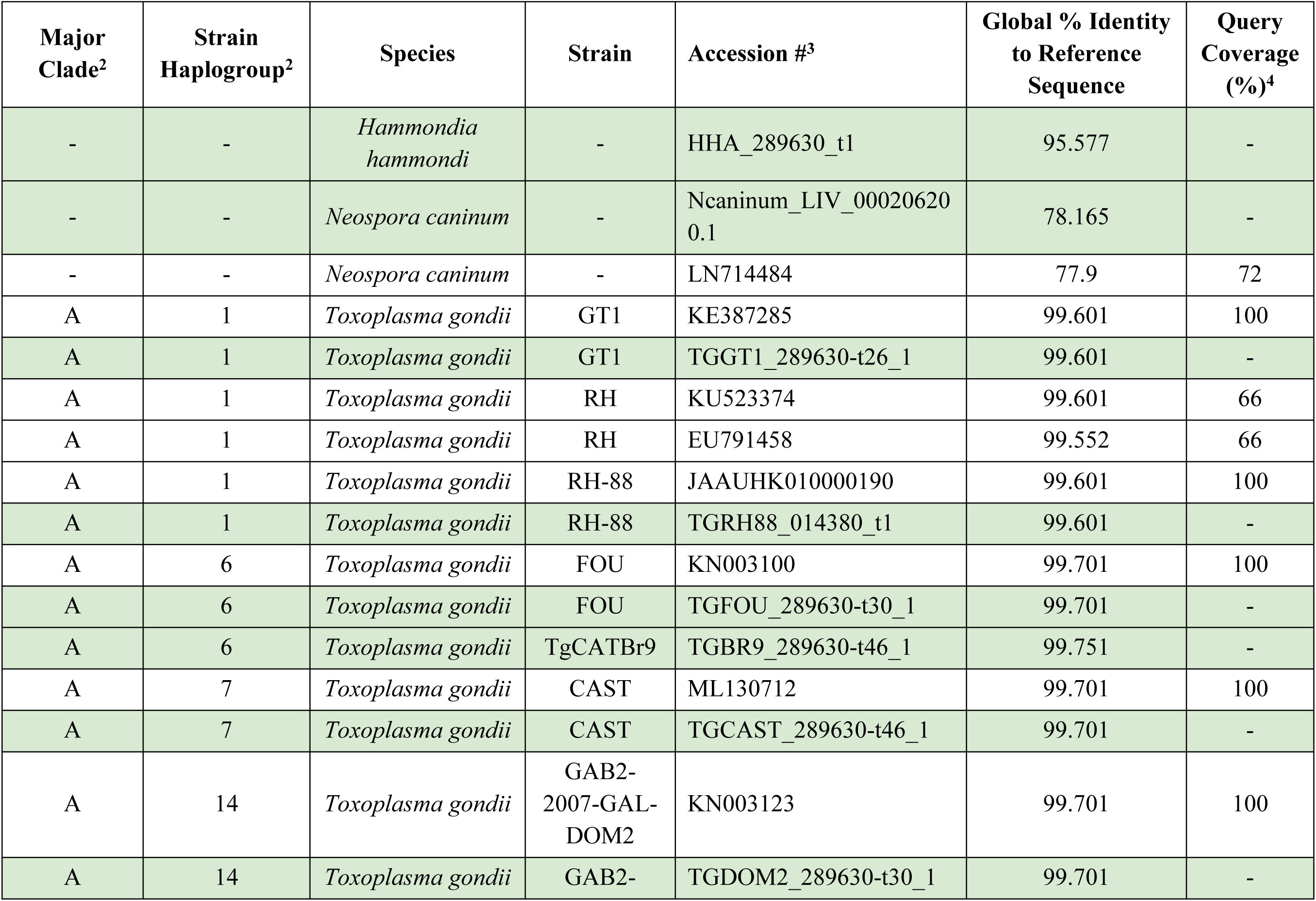
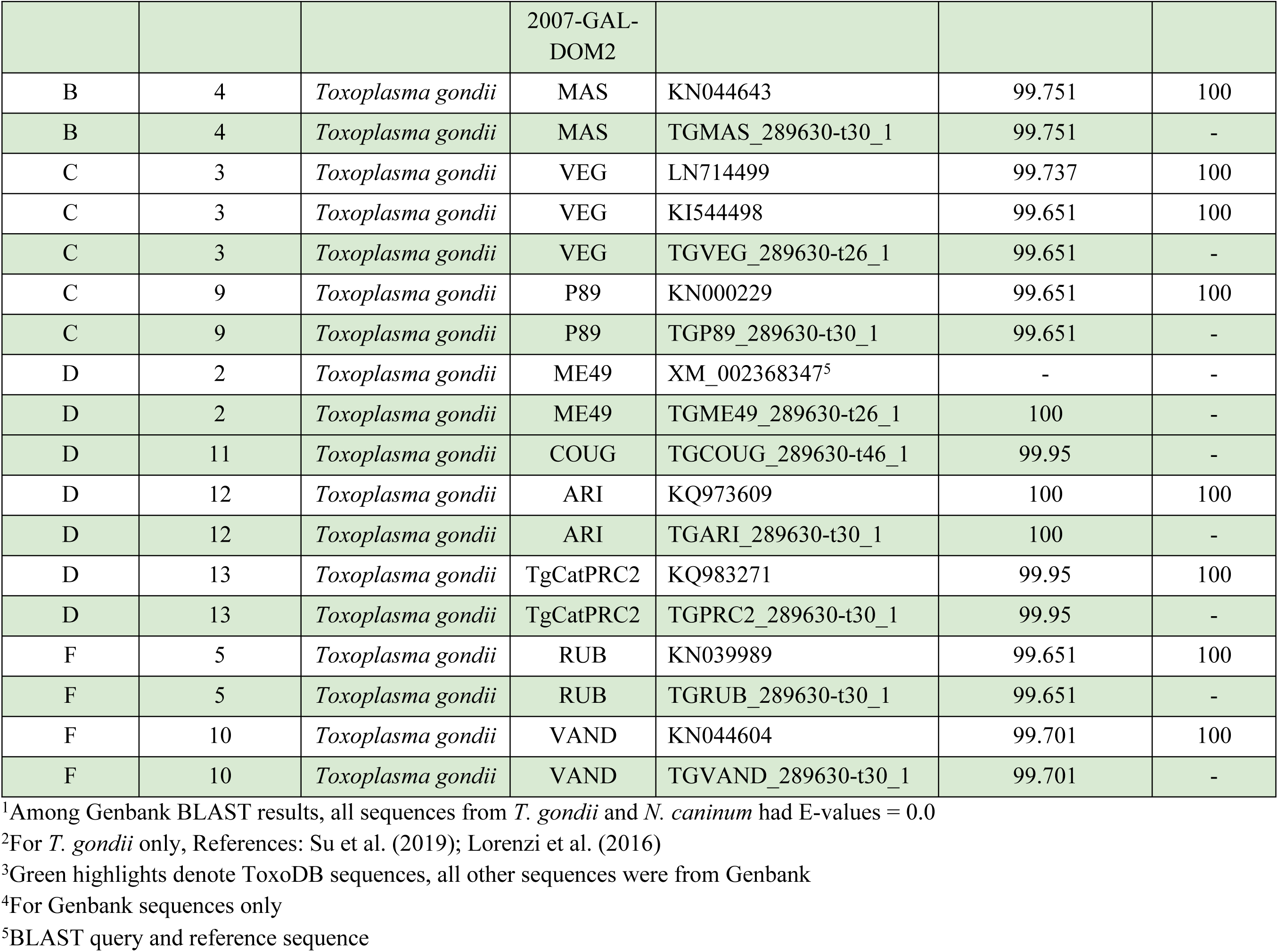
*MIC16* coding sequences from *Toxoplasma gondii* and outgroups.^1^.

The average CDS lengths of the three microneme protein genes were significantly different from each other (One-way ANOVA: F(2, 112) = 8815.54, p<0.0001, Tables 2-4). The average CDS length of *MIC12* was 4.5 times longer than that of *MIC13* and 3.2 times longer than that of *MIC16*, and the average CDS length of *MIC16* was 41.8% longer than that of *MIC13* (Tukey HSD post-hoc, p<0.01, Tables 2-4).

### Phylogenetic Analysis

Phylogenetic analysis was conducted to infer the evolutionary relationships between *T. gondii* strains for each of the three microneme proteins. Bootstrap values among the branches of the trees for the three MIC proteins vary widely. High coding sequence identity between *T. gondii* strains may contribute to some of the low bootstrap values (Figure 4). There is no significant difference between the average bootstrap values of cladograms constructed from *MIC13*, *MIC12*, and *MIC16* alignments (One-way ANOVA, F(2, 98) = 1.4, *p* = 0.2515, Figure 4).

**Figure 4.**
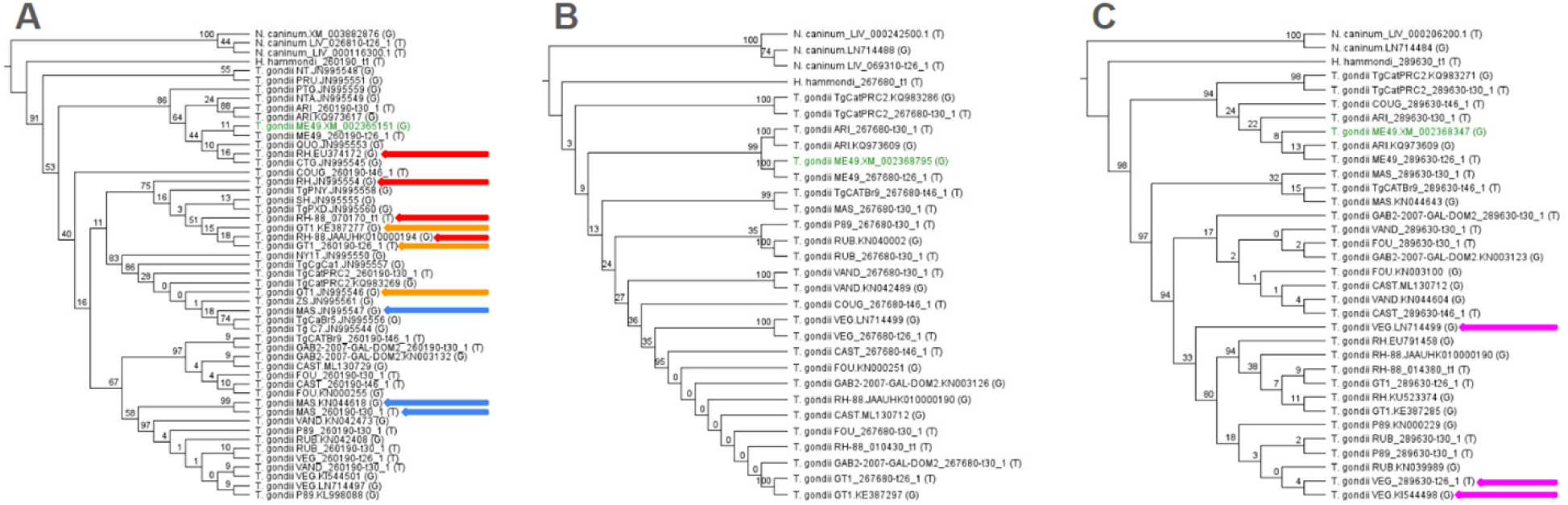
Maximum likelihood cladograms for *Toxoplasma gondii* strains based on *MIC13, MIC12,* and *MIC16* coding sequences. Tips are labeled with the species name and strain for *T. gondii* followed by Genbank accession number or gene number for ToxoDB sequences. *N. caninum* is set as the outgroup in all trees. Numbers on branches represent bootstrap values. Reference sequences are highlighted in green. When the same strain was sampled more than once and appears in different, well supported clades (bootstrap values ≥ 50), the polyphyletic strains have been noted with colored arrows.

We divided these trees into three parts to compare bootstrap support among sister relationships (recent splits), nodes with one tip (slightly deeper divergence), and internal nodes (including the most ancient relationships). For *MIC13*, there was no significant difference between the average bootstrap values for these three groupings within the ingroup (One-way ANOVA, F(2, 43) = 0.21, *p* = 0.8114, Figure 5). For *MIC12*, sister relationships had a 4.5 times higher average bootstrap value than non-sisters (One-way ANOVA, F(2, 22) = 24.53, *p* < 0.0001, Figure 5). Finally for *MIC16*, internal nodes had a 3.4 times higher average bootstrap value than sister relationships (One-way ANOVA, F(2, 27) = 4.07, *p* = 0.0285, Figure 5).

**Figure 5.**
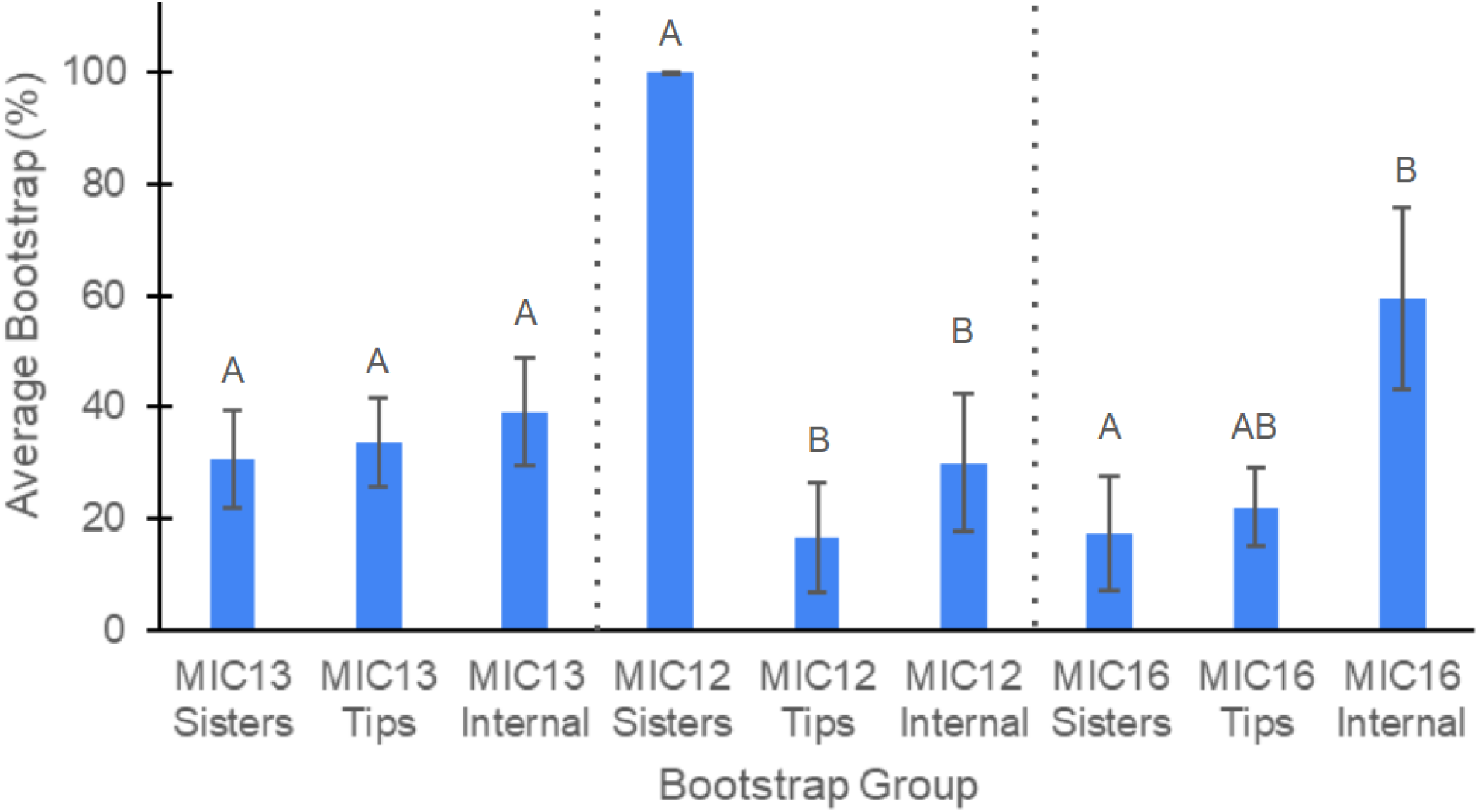
Comparison of average bootstrap values divided into three groups per gene based on coding sequences extracted from *Toxoplasma gondii* strains. Letters show Tukey HSD post-hoc test results within each MIC (separated by dotted lines); different letters within a MIC gene represent significant differences (*p* < 0.05). Nodes connecting sister groups are labeled sisters, nodes connected to one tip are labeled tips, and nodes connected to no tips are labeled internal. Error bars show standard error of the mean. Sample sizes: *MIC13* Sisters (n = 14), *MIC13* Tips (n = 19), *MIC13* Internal (n = 13), *MIC12* Sisters (n = 8), *MIC12* Tips (n = 10), *MIC12* Internal (n = 7), *MIC16* Sisters (n = 9), *MIC16* Tips (n = 13), *MIC16* Internal (n = 8).

Interestingly, some sequences listed under the same strain appear in different clades separated from each other by a bootstrap value of ≥ 50 (Figure 4). These include RH, GT1, and MAS in the *MIC13* tree and VEG in the *MIC16* tree (Figure 4). In each of these cases, the sequences from NCBI Datasets: Genome and ToxoDB remained in the same clade, while the other sequences from Genbank came from isolates which may have developed unique mutations.

Due to a higher number of strains represented in the sequence alignment for *MIC13*, we pruned the trees to include only overlapping strains for all three MICs to more easily compare the relationships between strains (Figure 6). There are 15 *T. gondii* strains shared across all three alignments. Due to low bootstrap values, a large number of nodes in each tree were collapsed into polytomies (bootstrap < 50%). The pruned trees show incongruent topology with strains that are strongly supported sisters in one tree being separated or part of a polytomy in another (Figure 6). Additionally, all three boxed colors can be traced to the same polytomy in the pruned trees for *MIC13* and *MIC12*, but in the pruned tree for *MIC16*, the green and blue boxed strains are more closely related to each other than to the blue boxed strains (Figure 6).

**Figure 6.**
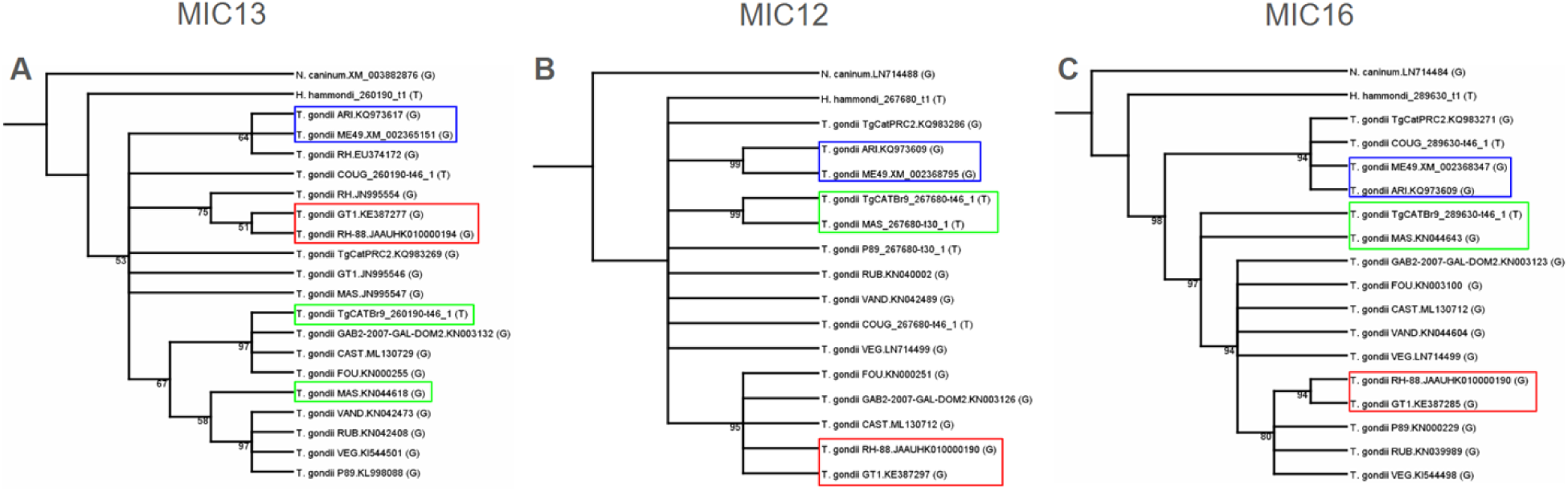
Pruned trees including only overlapping strains in all three MICs. Trees were pruned to just the strains present in all three individual MIC trees from Figure 6. Tips are labeled with the species and strain for *T. gondii* followed by Genbank accession number or gene number for ToxoDB sequences. *N. caninum* is set as the outgroup in all trees. Numbers next to nodes represent bootstrap values. Nodes with < 50% bootstrap support were collapsed into polytomies. For sister relationships between sequences from the same strain, only one sequence was kept. (A) *MIC13*, (B) *MIC12*, and (C) *MIC16*. If any of the three trees had a sister strain relationship, it received a colored box and then identified with the same colored box in the other two trees to determine consistency in sister relationships.

### Molecular Evolution

Positive selection was not detected by the SLAC algorithm in Datamonkey at any of the sites (representing codons) in any of the three MICs. SLAC detected purifying selection at one site in *MIC13* (in between the first sialic acid binding MARs from Genbank and the putative second sialic acid binding MARs from Ye et al., 2019) and 15 sites in *MIC12*, thus producing negative dN-dS values (Table 5). No signs of selection were detected in *MIC16* (Figure S1). The average dN-dS value across all sites in *MIC12* was 237 times higher than *MIC13* (dN-dS = 0.01 vs. −2.4) indicative of negative selection (Independent Sample *t*-test, t(2740) = 3.48, p<0.05, Figure 7). Additionally, the average dN-dS value across all sites in *MIC12* was negative, while that in *MIC13* was positive.

**Figure 7.**
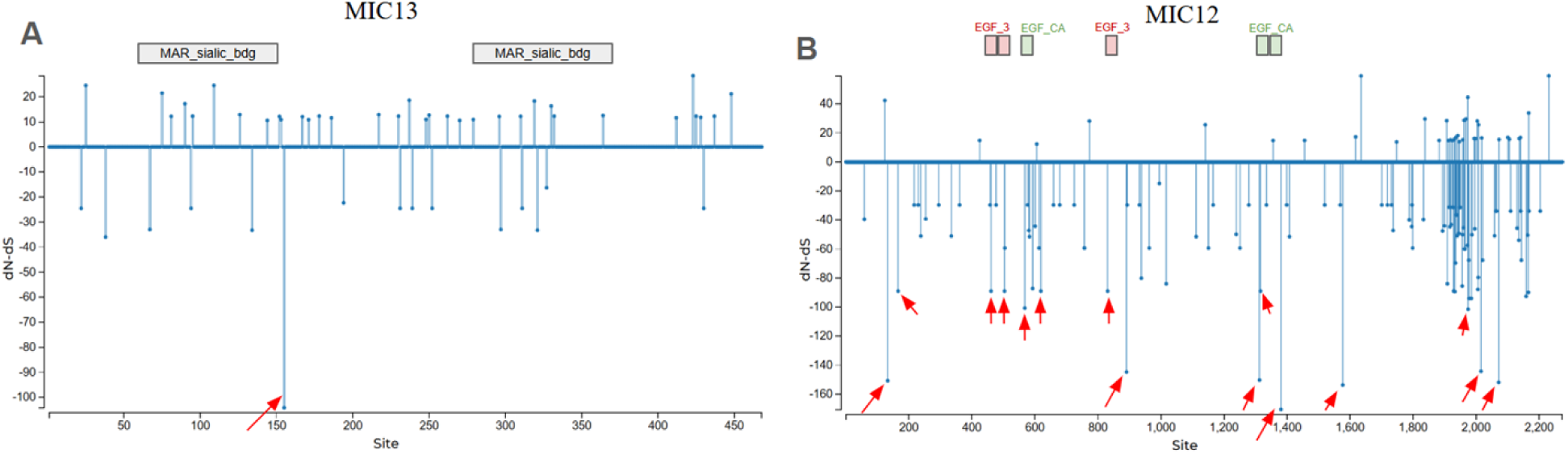
Difference in rates of non-synonymous and synonymous substitutions across codon sites for *MIC13* and *MIC12* coding sequences in *Toxoplasma gondii* strains. Differences within codons for amino acids were detected with SLAC. Sequences were compared across 468 sites in 51 sequences for *MIC13* (A) and 2274 sites in 30 sequences for *MIC12* (B). dN = nonsynonymous substitutions per nonsynonymous site. dS = synonymous substitutions per synonymous site. Red arrows indicate sites of significant purifying selection (*p* < 0.05). Boxed annotations represent MAR domains (A) and EGF domains that overlap with significant sites of selection (B, note: there are numerous EGF domains, but these are the only ones that overlap with significant purifying selected sites) as listed in Genbank protein flatfiles. MAR_sialic_bdg = Sialic acid binding micronemal adhesive repeats; EGF-3 = EGF domain; EGF-CA = Calcium-binding EGF-like domain.

**Table 5.**
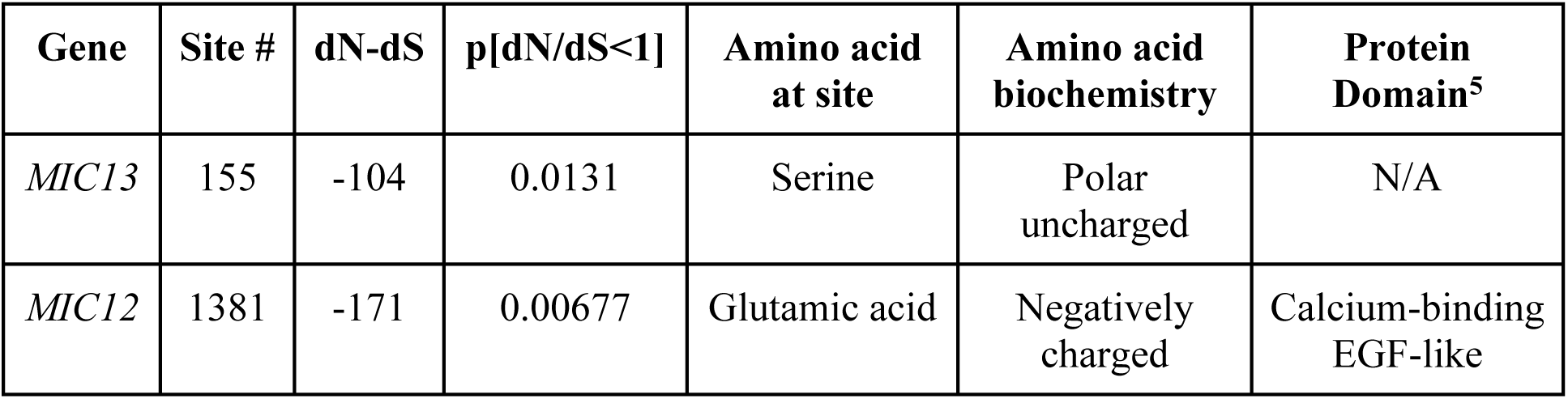
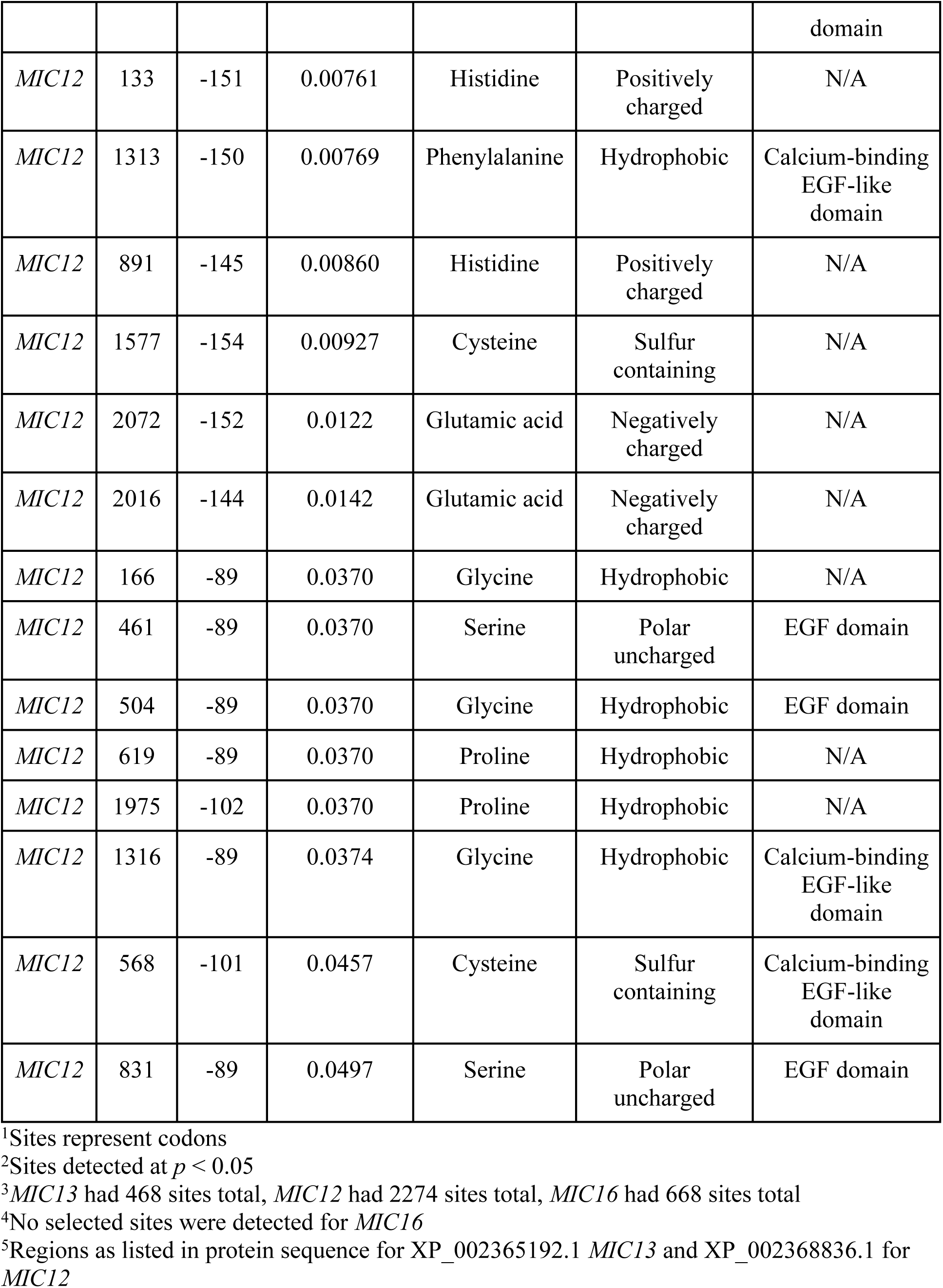
Sites^1^ of purifying selection in *MIC13* and *MIC12* of *Toxoplasma gondii* detected by SLAC.^2, 3, 4^.

### 3D Structures

There are no PDB structures available for MIC13, but an AlphaFold model of MIC13 was located (UniProt H9BC57). It has very low confidence for the first 50 amino acids, low confidence for the last three amino acids, and a few amino acids dispersed in the middle including E220, T264, T271, and G402, but otherwise high confidence in the 3D structure for the majority of the remaining amino acids (Figure 8). The predicted structure contains nine α-helices, 22 β-sheets, and 32 coil regions. There are two sialic acid-binding MARs from amino acids 57–150 and from amino acids 280–371. Neither of these regions coincides with site 155 where negative selection was detected (Table 5). Our 3D structure integration is limited to MIC13 since no empirical PDB structures were available and the AlphaFold models were highly uncertain for MIC12 and MIC16.

**Figure 8.**
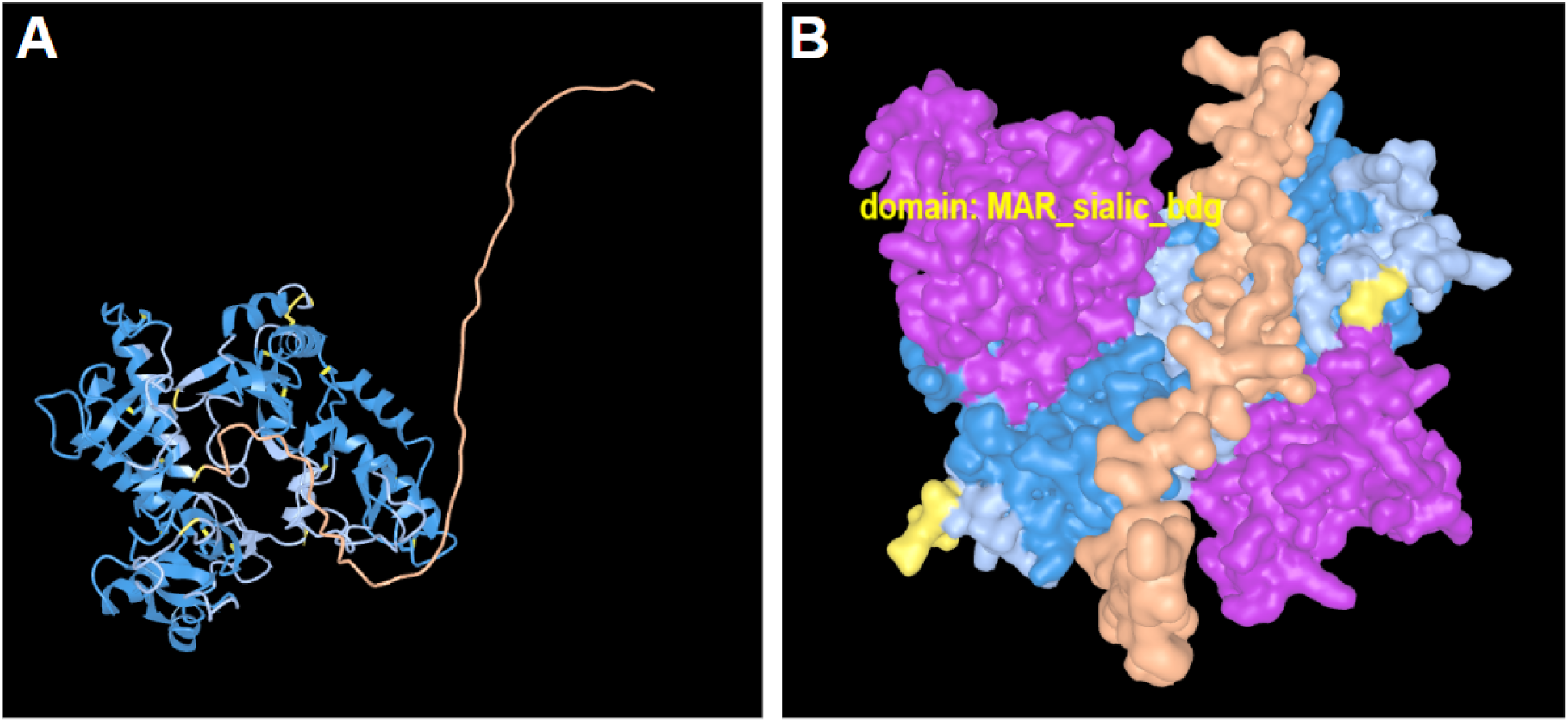
AlphaFold predicted 3D structure for the MIC13 protein (UniProt H9BC57) of Toxoplasma gondii. (A) Structure viewed in ICn3D and colored by confidence by pLDDT (dark blue = very high confidence, light blue = high confidence, yellow = low confidence, orange = very low confidence). (B) Same structure as (A) shown instead with van der Waals surfaces with sialic acid-binding MARs colored in purple.

## Discussion

In this study, we compared alignments, phylogenetic relationships, and molecular evolution in MIC13, MIC12, and MIC16 from *T. gondii*. Although all three proteins are localized in the micronemes, they did not show the predicted pattern of rapid adaptive evolution. Instead, the three genes showed discordant phylogenetic histories, no evidence of positive selection, and heterogeneous levels of purifying selection, with strongest constraint in MIC12. Below, we interpret these findings in light of previous phylogenetic, genomic, and functional studies of *T. gondii* MIC proteins.

### Phylogenetic Analysis

Phylogenetic analyses of the three MIC coding sequences indicate that these loci do not recover a single shared strain history, consistent with previous genome-scale studies of *T. gondii* population structure and admixture (Grigg and Sundar, 2009; Minot et al., 2012; Lorenzi et al., 2016). Among the three genes, MIC12 generally showed stronger support for several recent sister relationships, whereas MIC16 showed somewhat stronger support at deeper internal nodes, suggesting that phylogenetic signal may be distributed differently across these loci. Using largely the same GenBank sequences as Ren et al. (2012), we also broadly recovered their MIC13 relationships while adding additional representation of VEG and ME49 strains.

We recovered several strain relationships consistent with whole-genome phylogenetic analyses of *T. gondii* (Lorenzi et al., 2016). For example, the close relationship between the reference strain ME49 and ARI (haplogroup 12) observed across all three MICs aligns with Lorenzi et al.’s report that these strains share ∼60% conserved haploblocks. Similarly, in the MIC16 tree pruned to overlapping strains (Figure 6C), we identified a strongly supported clade containing TgCtPRC2 (haplogroup 13), COUG (haplogroup 11), and ME49. Lorenzi et al. (2016) whole-genome comparisons report that TgCtPRC2 and ME49 share ∼40% conserved genomic regions, whereas TgCtPRC2 and COUG only share less than 1% and are less likely to be as closely related as our MIC16 results suggest - discordance between gene-level and genome-wide relationships. Additional inconsistencies were evident for strains with proposed recent origins. For example, although Minot et al. (2012) suggest that GT1 arose from a ME49 × BOF cross, our GT1 sequences consistently group with RH-88 (as expected since these are both Type I strains) and remain distinct from ME49 across all three MICs (Figure 6; note we did not sample BOF). In MIC13, GT1 appears in multiple clades (Figure 4, all samples tree, orange arrows), whereas in MIC12 it forms a strongly supported sister relationship and shows no polyphyly in MIC16. Similarly, VEG, which Minot et al. (2012) propose originated from a ME49 × P89 cross, does not group with ME49 in any of our three trees pruned to the overlapping strains (Figure 6; note we did not sample P89). In the full dataset (Figure 4), two VEG sequences are closely related in MIC13, strongly supported sisters in MIC12, and strongly polyphyletic in MIC16. Together, these examples demonstrate that individual MIC loci recover discordant strain relationships that do not consistently reflect genome-wide expectations.

These discordances indicate that MIC loci do not track a single bifurcating strain history in *T. gondii*, consistent with prior genome-scale studies showing admixture, recombination, and mosaic genome structure (Minot et al., 2012; Lorenzi et al., 2016). Because *T. gondii* undergoes sexual recombination within and among strains, different loci may retain distinct evolutionary histories shaped by recombination and the inheritance of large conserved haploblocks (Minot et al., 2012; Lorenzi et al., 2016). Selective sweeps of advantageous gene combinations within these haploblocks may further decouple gene genealogies from genome-wide phylogenetic patterns. At the same time, rare recombination events within otherwise clonal populations may obscure phylogenetic signal and complicate reconstruction of strictly bifurcating relationships. Consequently, the incongruence observed among MIC trees is consistent with a combination of admixture, incomplete lineage sorting, and locus-specific evolutionary processes, although the present data do not distinguish among these alternatives. Determining whether these MICs reside within conserved haploblocks or the more variable intervening genomic regions will be important for understanding the evolutionary forces shaping their histories and patterns of constraint.

### Molecular Evolution

Molecular evolution analyses reveal varying degrees of evolutionary constraint within individual MIC alignments and among the three MIC proteins. Across all three genes, no sites exhibited evidence of positive selection and relatively few sites showed significant purifying selection, with all but one of these sites occurring in MIC12. In MIC12, 15 sites exhibited significant site-specific purifying selection, accompanied by an overall negative average dN–dS, indicating strong evolutionary constraint. In contrast, MIC13 contained only a single site under purifying selection, and no significant selection was detected in MIC16. These results suggest that MIC12 is under stronger functional constraint, whereas MIC13 and MIC16 may experience weaker or more context-dependent constraint (or most sites are relatively neutral). For MIC12, many of the sites under purifying selection overlap with EGF-Ca and EGF-3 motifs, extracellular structural domains that stabilize adhesion interfaces and depend on conserved cysteine residues and, in some cases, calcium-binding sites. The strong purifying selection observed in these regions likely reflects stringent structural requirements for host-cell binding and invasion which fits with some experimental work on MIC16 showing that it is not associated with host cell infection, but instead required for the optimal growth of parasites under stress or bradyzoite-inducing conditions (Ye et al., 2019).

These findings contrast with previous suggestions of positive selection in MIC16 (Liu et al., 2016), although that study did not report codon-based dN/dS estimates, limiting direct comparison. Instead, our results are consistent with genome-wide analyses of molecular evolution in *T. gondii* (Minot et al., 2012; Lorenzi et al., 2016), which found little evidence of positive selection across most genes. For example, Minot et al. (2012) did not identify any MICs among the top 18 genes with the highest numbers of non-synonymous substitutions across 45 genomes (their Table S4). Similarly, after reviewing the Supplemental Data provided by Lorenzi et al. (2016), we found relatively low dN/dS values for our target MICs (MIC13 = 0.58; MIC16 = 0.528; MIC12 = 0.079), with MIC12 ranking among the most strongly constrained MICs in their dataset. In contrast, genes with strong signatures of positive selection are more commonly found in other families, like the dense granule proteins (GRA; dN/dS > 5, also mentioned in Hakimi, 2022) and SAG-related sequences (SRS; dN/dS > 3) identified by Lorenzi et al. (2016) and the concentration of rhoptry organelle proteins in the 18 genes with the most non-synonymous substitutions (ROP1, ROP16, ROP17, ROP18, ROP39) as suggested by Minot et al. (2012). Together, these comparisons indicate that, despite the putative roles of these three MICs may have in host cell invasion (and other developmental functions per Ye et al., 2019), they are not primary targets of rapid adaptive evolution (nor are any of the other microneme proteins including AMA1 dN/dS = 0.42 per Lorenzi et al, 2016 Supplemental Data), but instead appear to be evolutionarily constrained by essential structural and functional requirements.

### 3D Structure of MIC13

The predicted 3D structure of the MIC13 protein includes two sialic acid-binding micronemal adhesive repeat (MAR) domains positioned on either side of the protein, similar to those described in MIC1 (Friedrich et al., 2010). Because sialic acid residues are abundant on the surface of mammalian cells, these MAR domains are thought to facilitate host-cell adhesion and invasion. This provides a structural context for interpreting the limited evidence of selection in MIC13, suggesting that conserved adhesion functions may constrain sequence evolution.

However, functional evidence for MIC13 remains ambiguous. Ye et al. (2019) reported no significant difference in invasion efficiency between MIC13-disrupted mutants and parental strains, indicating that MIC13 may be functionally redundant or that its role depends on the developmental stage and context in which it is expressed. This functional uncertainty is mirrored by structural ambiguity. While the AlphaFold model used here predicts two MAR domains, previous work inferred three such domains (Ye et al., 2019), highlighting uncertainty in both the number and positioning of these motifs. The absence of experimentally determined structures further limits precise mapping of functional regions.

Taken together, the predicted presence of adhesion-related MAR domains alongside limited evidence of purifying selection and no detectable positive selection suggests that MIC13 may operate under moderate structural constraint without being a primary target of diversifying selection. Resolving the functional importance and structural organization of these domains will require experimental validation of MIC13 structure and function.

### Limitations and Future Directions

The sequence dataset used in this study was derived primarily from publicly available genomes in NCBI, with relatively few additional isolates retrieved through targeted BLAST searches. As a result, sampling may be biased toward well-characterized laboratory and reference strains. In addition, MIC16 sequences reported by Liu et al. (2016) (GenBank accessions KX147274–KX147285) were not present in the core_nt database and could not be independently verified, and therefore were not included in our analyses. This limited our ability to fully replicate their phylogenetic and molecular evolution results.

A second limitation involves uncertainty in protein annotation and domain prediction. Molecular evolution analyses relied on features annotated in GenBank, which may not reflect current or experimentally validated domain structures. For example, the number and positions of sialic acid-binding MAR domains in MIC13 differ among studies (Genbank vs. Ye et al., 2019), complicating interpretation of structure–function relationships. In addition, a “PspC subgroup 2” domain was annotated in the MIC12 reference sequence despite PspC being a pneumococcal surface protein from *Streptococcus pneumoniae*, and the corresponding region showed no meaningful similarity to bacterial sequences (XP_002368836.1). This annotation was therefore excluded from our analyses, highlighting the need for careful curation of predicted protein features.

Finally, none of the MIC proteins examined here have experimentally determined 3D structures. Structural analyses of MIC13 relied on AlphaFold predictions, which showed low confidence in the N-terminal region and differed from previous studies in the number of predicted MAR domains. To assess structural consistency, we compared multiple AlphaFold models from *T. gondii* and related apicomplexans using VAST, which supported overall structural similarity across species, including with *N. caninum* (Figure S2). However, predicted structures remain an imperfect proxy for experimentally resolved models, and future structural and functional studies will be necessary to further validate domain organization and refine interpretations of evolutionary constraint.

## Conclusion

These three MICs did not show the predicted pattern of rapid adaptive evolution. Instead, they showed high sequence conservation, discordant gene trees, and strongest constraint in MIC12, especially in EGF-related domains. This suggests that at least some microneme proteins are constrained by essential structural or functional requirements rather than shaped primarily by recurrent host–parasite arms races.

